# Mitigating consecutive drought impacts on forest productivity through strategic tree species spatial design

**DOI:** 10.64898/2026.04.13.718112

**Authors:** Wentao Yu, Ulrich Brose, Benoit Gauzens

## Abstract

The rising frequency and severity of multiyear droughts due to climate change poses a serious threat to tree growth and survival, compromising our terrestrial carbon sink. Although tree diversity is known to enhance forest biomass production, its role in mediating drought impacts remains elusive due to inconsistent evidence and limited understanding of species interactions. Using empirically parameterized pairwise interactions among eight tree species, we show that tree diversity buffers forest communities against repeated drought-induced biomass loss. This buffering arises from heterogeneous neighborhoods that promote both niche differentiation and facilitative interactions among species. We further demonstrate that strategic planting designs, such as random or single-line spatial arrangements, amplify these benefits by maximizing neighborhood heterogeneity, with single-line being more plausible when balancing management effort. Our simulation results suggest that increasing diversity could raise carbon sequestration rates by 18.8%. These findings corroborate tree diversity and spatial heterogeneity as actionable, climate-adaptive tools that simultaneously enhance forest productivity, drought resilience, and long-term carbon sequestration.

## Introduction

Climate change is intensifying due to rising greenhouse gas emissions, driving long-term warming trends that are closely associated with an increase in climate extremes (Diffenbaugh et al., 2017). A key mitigation strategy for climate change is forest expansion and management, as forests can accumulate substantial carbon in tree biomass over time (Griscom et al.,2017; Kauppi et al., 2022). Over recent decades, global forests have been acting as a steady carbon sink of about 3.6 ± 0.4 petagram Carbon year^−1^ in the 1990s-2000s, and ∼3.5 ± 0.4 petagram Carbon year^−1^ in the 2010s (Pan et al. 2024). Within this context, mixed-species forests are particularly promising: diverse tree communities typically achieve higher productivity and carbon storage (Duffy, Godwin, & Cardinale, 2017; Huang et al., 2018; Liu et al., 2018), which indirectly buffers against climate change. Tree diversity thus reduces risks that climate stress shifts ecosystems from carbon sinks to sources that accelerate climate change (Zhang at al., 2025). These positive effects of tree biodiversity on biomass production are driven by ecological mechanisms such as the selection of dominant producers, niche complementarity of tree species and systematic patterns in pairwise tree interactions (Loreau & Hector 2001; Albert et al., 2022; Yu et al 2024), which need to be harnessed to maximize carbon sequestration.

Despite their critical role in mitigating climate change, forests remain highly vulnerable to its impacts. A major consequence from climate change is the rising frequency and severity of multiyear droughts, as demonstrated by the record-breaking 2018–2020 Central European drought (Hari et al. 2020; Rakovec et al. 2022). According to the insurance hypothesis (Yachi, S., & Loreau, 1999; Loreau 2021), a mixture of tree species with contrasting climatic optima in their growth rates should help to maintain the functioning of forests under highly variable climatic conditions. Indeed, mixed forests have demonstrated higher temporal stability of productivity under fluctuating environments (Schnabel et al.,2019, 2021), suggesting that tree diversity could buffer against environmental variability. However, the role of tree diversity under extreme drought stress is less clear. While drought is generally known to suppress tree growth (Allen et al., 2015; Bouwman et al., 2025), it remains poorly understood how biodiversity interacts with drought to shape post-drought recovery and long-term biomass production. Previous studies report that the effects of tree diversity under drought range from positive to neutral or even negative under drought of varying intensity (Grossiord, 2014; Forrester et al., 2016; Grossiord, 2020), suggesting that increasing tree species diversity alone might not necessarily improve trees’ ability to face increasing drought stress (Serrano-León 2025). Consecutive drought years, in particular, can exacerbate initial impacts through cumulative soil (Anderegg et al., 2020; Schnabel et al., 2022), with potentially severe consequences for forest plantations. Given the increasing frequency and intensity of multiyear drought events globally, there is a large knowledge gap about the efficiency of forest management strategies in face of these events.

Beyond species diversity, the spatial design of trees represents a crucial yet often overlooked component of forest management strategies. Pairwise interactions between neighboring trees critically influence individual tree growth, with interspecific interaction being systematically less negative, or even positive, compared to intraspecific interactions (Yu et al. 2024). These effects scale up to determine how tree diversity drives community-level productivity and carbon sequestration (Yu et al. 2024). In the following, we will refer to the effect of this specific structure in the strengths of inter-compared to intra-specific interactions on tree productivity as the *interaction structure effect*. Because the spatial arrangement of trees determines how often species interact with conspecifics, leading to strong competition, or other tree species, yielding comparatively weaker competition or facilitation, the spatial design of planting trees can substantially enhance ecosystem functioning by optimizing the interaction distribution in the local neighborhood (i.e., increasing heterogeneity; Fig.1 A). In line with this theoretical prediction, recent evidence demonstrates that promoting the spatial heterogeneity in tree species planting can significantly improve multiple ecosystem functions, including biomass production, litterfall distribution, and litter decomposition rates (Supplement S1,2; Beugnon et al., 2025).

**Fig. 1.**
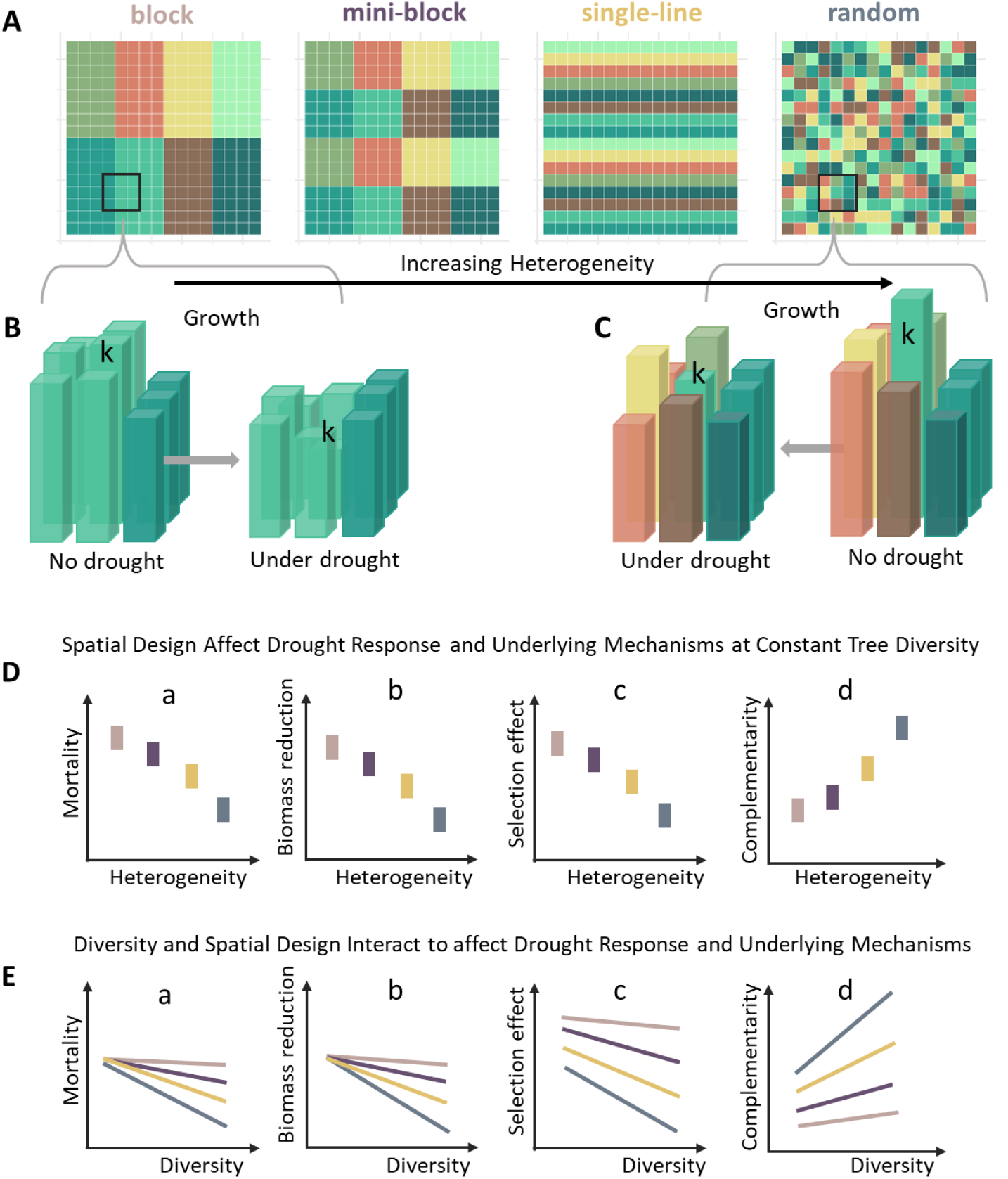
Conceptual illustration of how spatial design and diversity interact to affect drought response in forest plantations. (A) Four spatial designs for an eight-species mixture with increasing spatial heterogeneity. Each grid cell represents a tree, colored by species identity, and each forest stand comprises 16 × 16 trees. Insets show neighborhood compositions in block and random designs. (B) A homogeneous neighborhood from the block design and (C) A highly diverse neighborhood from the random design, showing growth of trees in these two neighborhoods under normal conditions versus consecutive drought events. (D) Hypothesized drought responses across the four spatial designs at constant diversity. (E) Hypothesized drought responses across the four spatial designs at varying diversity levels.

In contrast to these effects on tree productivity, little is known about how the spatial planting design interacts with tree diversity to drive forest responses to drought. Extreme drought events impose strong stress on trees by limiting water and nutrient availability, thereby reducing growth, and increasing mortality risk (Breshears et al., 2005; Allen et al., 2015). Because tree species differ in their ability to tolerate and recover from drought, their growth performance varies before, during and after drought events (Hesse et al., 2023; Tao et al 2024). Drought can thus shift selection effects from being driven by species with the fastest growth to those with the highest drought tolerance or quickest recovery. These shifts can be further modified by the niche complementarity of neighboring species, as differences in abiotic niches facilitate recovery through non-overlapping resource use. Similarly, weaker competition or facilitation in the local neighborhood can also enhance recovery or mitigate drought impacts.

For example, in Fig. 1B the focal species k (center) is a fast-growing dominant species more susceptible to strong growth reductions under drought (McGregor et al., 2016; Guillemot et al., 2022). In this homogeneous scenario, the focal tree is surrounded mostly by conspecifics. Drought conditions could weaken the focal tree’s competitive ability, exposing it to intense competitive pressure from its homogeneous neighborhood and potentially leading to mortality (Fig.1 D-a and -b). Consequently, selection effects will be reduced (Fig.1 D-c). Conversely, in the heterogeneous scenario (Fig.1 C), the focal tree is surrounded by neighbors with diverse ecological strategies (e.g., drought tolerance, rooting depth, water-use efficiency) and weaker competition. Such heterogeneous neighborhoods greatly foster differential resource use and facilitative interactions under stress environments (Bertness & Callaway, 1994; Holmgren& Scheffer 2010), thereby buffering drought-induced biomass loss or mortality through amplifying effects of niche complementarity and weakened competition by other species (Fig.1D-d). Thus, spatial heterogeneity is expected to modulate drought-induced biomass reduction by altering interaction structure effects and shifting the relative importance of selection and complementarity. Designing forests with optimized spatial heterogeneity therefore offers a potentially powerful strategy to make mixed-species plantings not only more productive but also more resilient in maintaining productivity under consecutive drought events. Yet, empirical tests on the importance of tree interactions for facing consecutive droughts remain elusive.

Leveraging a tree growth model parameterized by a large-scale tree biodiversity-ecosystem functioning (BEF) experiment in subtropical China, we conducted a simulation experiment in a full-factorial design across a gradient of tree species richness (one, two, four and eight species) and four spatial planting designs: blocks, mini-blocks, single-line, and random (see Fig. 1 A for examples). We simulated three consecutive drought events by reducing species-specific growth rates. The species-specific reduction in growth rates were informed by empirically measured drought-related traits drought-related traits (see method for detail). To capture variations in post-drought recovery, we assigned species-specific recovery rates based on their fast-slow growth strategies (see method for detail). With this framework, we addressed three key questions: (1) Which spatial design most effectively maintains productivity under consecutive drought events, and does this design also maximize productivity under normal conditions? (2) How does diversity interact with spatial design to affect tree species’ response to consecutive drought events (Fig.1 E)? (3) Through which mechanisms do spatial design and diversity mitigate biomass loss under consecutive drought events?

## Results

We simulated tree community biomass growth under drought using empirical estimates of species-specific growth rates, pairwise intra- and interspecific interaction strengths, and drought tolerance. Simulations were conducted in a full-factorial design across a gradient of tree species richness (one to eight species) and four spatial planting designs: random, blocks, mini-blocks, and lines (Fig. 1 A). Each of these simulations were compared to the outcome of simulations with the same interaction and spatial configurations parametrization but without growth reduction induced by drought events.

### Total biomass reduction and tree mortality across diversity and spatial designs

Across all four spatial designs, the proportion of total biomass reduction caused by consecutive drought events consistently declined with increasing diversity (Fig. 2A). Consequently, the more tree species coexist in a community, the lower the biomass loss following a drought. Among the four spatial designs, random and single-line arrangements buffered drought effects more effectively across diversity levels, experiencing markedly lower biomass reduction than block and mini-block designs, particularly at higher diversity levels. On plots with two tree species, the single-line design exhibited a slightly lower proportion of biomass loss than the random design. This likely reflects higher species heterogeneity in two species single-line arrangements (Beugnon et al. 2025), which translates into more heterogeneous interaction distributions in the local neighborhood than random designs that, at these low levels of diversity, may form single-species clusters, thereby decreasing the spatial heterogeneity of neighborhoods. On plots with eight tree species, however, random designs generally result in higher neighborhood heterogeneity (see Fig. 1A, C) and thus show lower biomass loss than single-line arrangements. Specifically, eight-species mixtures with random design yielded 373 ± 33 kilograms (mean ± standard deviation) more biomass than the block design (Supplement S3: table.1), more effectively buffering the detrimental effects of drought. Together, these results suggest that tree biodiversity buffers forest communities against drought-induced biomass loss, and that this buffering effect is enhanced when trees are planted in random or single-line designs.

**Fig. 2.**
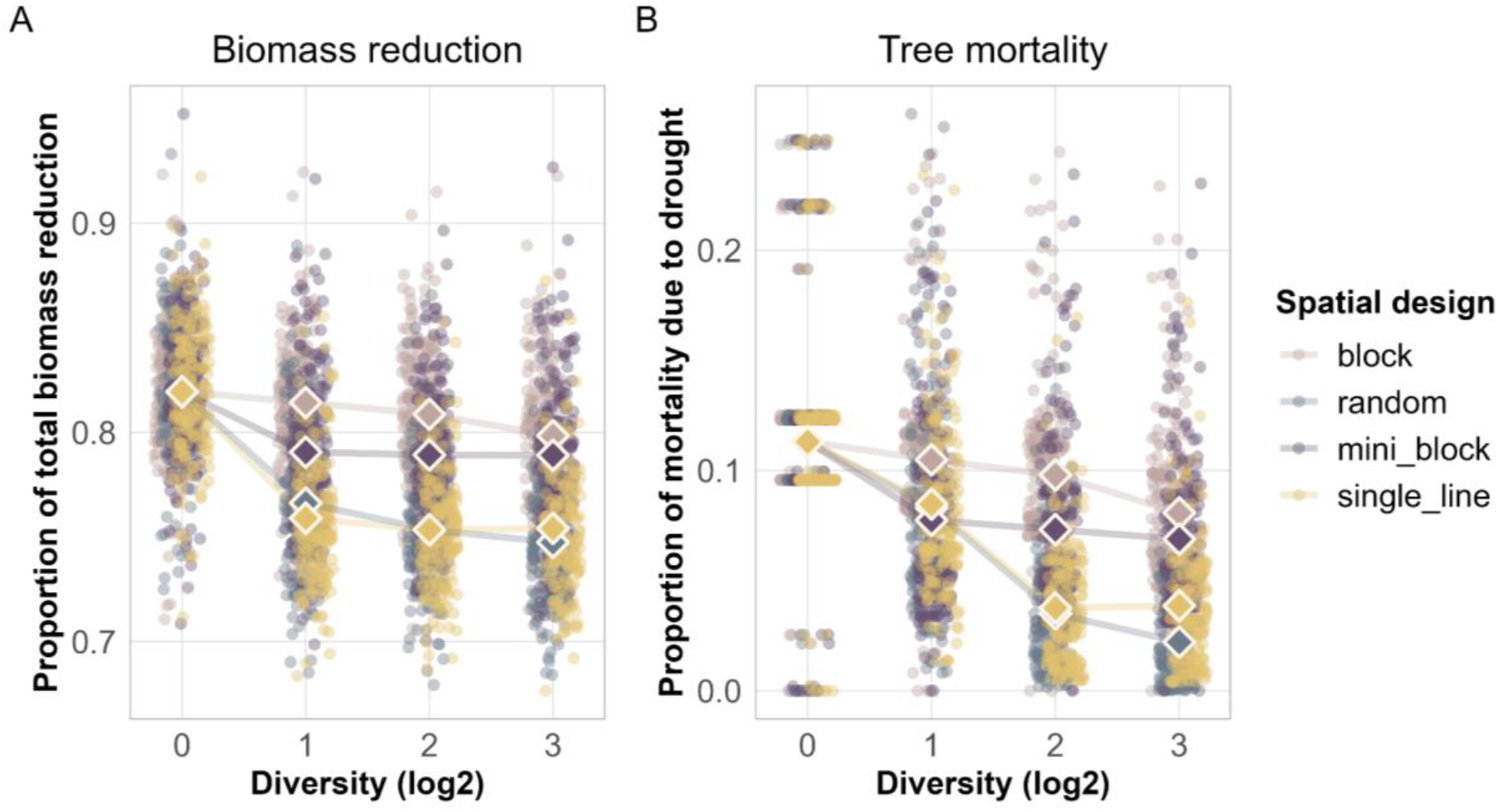
Effects of consecutive drought events on tree community biomass and mortality across diversity gradients and spatial designs (block, mini_block, single_line, random). (A) Proportion of total biomass reduction and (B) proportion of tree mortality in different spatial designs along a diversity gradient. Individual points represent plot-level values (N = 3,200), while squares indicate mean values for each diversity level within respective spatial designs.

**Fig. 3.**
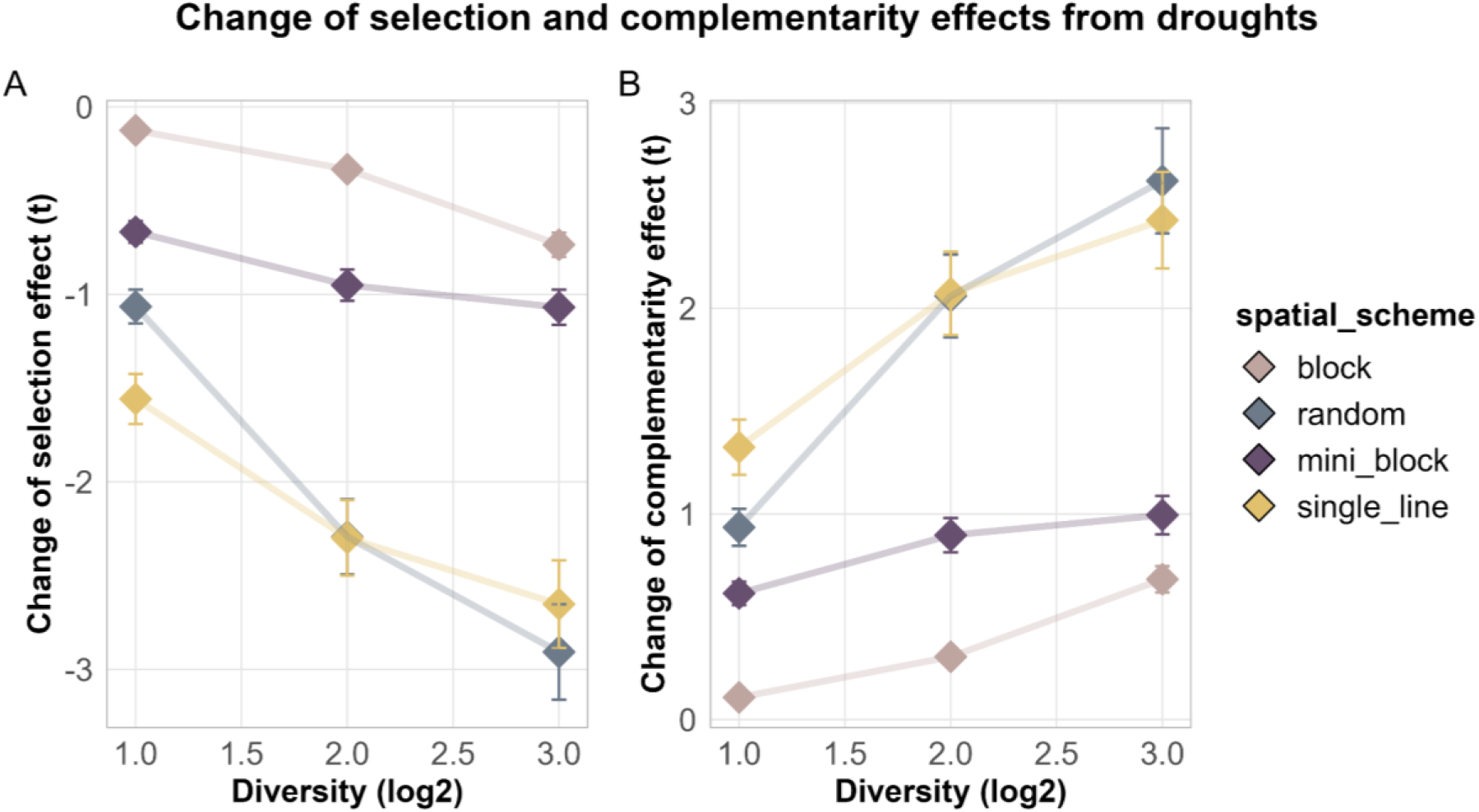
The change of selection (A) and complementarity (B) effects induced by consecutive drought events across diversity gradients and spatial designs. Squares denote mean values of selection and complementarity effects across all plots at each diversity level within a given spatial design, respectively, and error bars indicate standard errors.

Similar to biomass reduction, tree mortality from consecutive droughts declined with increasing diversity across all four spatial designs. At four- and eight-species diversity levels, single-line and random design provided the strongest buffering effect, with considerably lower mortality rates compared to block and mini-block designs. The parallel patterns for biomass reduction and tree mortality indicate that a more heterogeneous local neighborhood in terms of species composition can better buffer drought impacts by reducing tree mortality.

### Tree-tree interaction structure mediating biomass responses to consecutive drought events

#### Community-level mechanisms driving biomass responses to consecutive drought events

By partitioning net diversity effects into selection and complementarity effects under drought and non-drought conditions, we examined how drought events modify the mechanisms driving net diversity effects across tree diversity gradients and spatial designs. Overall, drought weakened selection effects, which also diminish as diversity increases across all four spatial designs. The steepest declines occurred in random and single-line configurations, suggesting that repeated drought stress disproportionally suppresses dominant, high-productivity species. Notably, this suppression is stronger in more diverse communities.

In contrast, complementarity effects strengthened with increasing diversity across all spatial designs, with the strongest gains observed in random and single-line arrangements. This finding indicates that under severe drought conditions, facilitative and niche differentiation processes become increasingly more important for maintaining productivity. The spatial design of tree species strongly influenced these effects: block and mini-block designs exhibited markedly weaker productivity increases than random and single-line arrangements at comparable diversity levels. These findings imply that spatial clustering limits opportunities for niche complementarity or facilitation among species, whereas heterogeneous neighborhoods, where trees with diverse drought-tolerance, growth strategies and interaction strengths are intermixed, enhance complementarity and interaction effects, especially during drought events.

Overall, the relative importance of underlying mechanisms was shifted substantially by droughts: in more diverse communities with heterogeneous neighborhoods (i.e., random and single-line designs), complementarity effects became the primary driver of net diversity effects under drought. This shift likely underpins the considerably lower mortality observed in the diverse communities with random and single-line design.

## Discussion

Drought-induced declines in tree growth and survival are increasing globally with climate change, threatening the terrestrial carbon sink (Anderegg et al. 2020). While ample evidence supports the beneficial effects of tree diversity on forest biomass production (Liang et al., 2016; Feng at al., 2022), the role of tree diversity in mediating drought impacts remains unclear due to inconsistent evidence and poor understanding of underlying processes, particularly how species interactions mediate drought impacts on individual trees. Using empirically parameterized pairwise interactions among eight tree species, we show that tree diversity can buffer forest communities against repeated drought-induced biomass loss. This buffering emerges from the formation of heterogeneous neighborhoods that promote both niche differentiation and facilitative interactions amongst different tree species. Importantly, we demonstrate that this buffering effect can be amplified through strategic planting to maximize neighborhood heterogeneity by random or single-line spatial design. Our findings therefore not only illuminate the mechanisms by which diversity buffers drought impact, but also establish spatial design as an effective, actionable tool for climate-adaptive forest management.

### Diversity and spatial design jointly affect drought response

Positive diversity effects on ecosystem functioning can stem from selection of dominant producers, complementarity effects reflecting niche differentiation and facilitation, and the distribution of interaction strengths (Loreau & Hector 2001; Barry et al., 2019; Yu et al., 2024). Because interactions at the individual scale aggregate to shape ecosystem functioning at the community scale (Fichtner et al., 2018; Yu et al., 2024), the capacity of diversity to mitigate drought impacts on tree growth ultimately depends on interactions among local neighbors. Our results reveal that increasing diversity fosters more heterogeneous neighborhoods, in which each tree individual has higher chances of being surrounded by other tree species. This increases the relative frequency of interspecific interactions over intraspecific interactions, which are, on average, less negative than intraspecific interactions (Yu et al. 2024). Our results reveal that this interaction structure in heterogeneous neighborhoods reduces biomass loss under consecutive drought events. This aligns with previous findings that neighborhood tree species richness can buffer drought-induced growth decline (Fichtner et al., 2020). We further show that higher diversity consistently lower drought-induced mortality, with the strongest reductions observed in the most diverse stands (eight species). Unlike declines in growth, which trees can recover from, tree mortality represents a permanent loss of carbon storage and can trigger cascading declines in various ecosystem functions. Together, these results stress the importance of tree diversity for forest drought resistance and suggest shifts in the realized interaction strength distribution when comparing homogeneous monocultures with heterogeneous forests of higher diversity as a mechanistic explanation.

Besides these beneficial effects of tree diversity, our results demonstrate that even at a given level of diversity, the resilience of forest communities can be substantially enhanced through strategic spatial design of forest stands. Maximizing the heterogeneity of local neighborhoods significantly amplifies the buffering effect arising from tree interactions against drought, with random spatial arrangements providing the strongest protection for both growth and survival in eight-species mixtures, followed closely by single-line designs. This result agrees with Beugnon (2025), who found that tree biomass yield under normal conditions was highest in random designs, with single-line designs ranking second. Interestingly, our results reveal that both tree diversity and spatial planting design enhance drought resistance by increasing neighborhood heterogeneity, which in turn shifts species interactions toward weaker competition. Tree diversity and spatial design used in forest management has the potential to enhance multiple ecosystem functions, including productivity and litter decomposition (Beugnon et al. 2025). Our results extend these recent findings by revealing how these two factors jointly shape drought response, enhancing resilience and sustaining ecosystem functioning under climate change.

### Shifts in selection and complementarity effect under drought events

Traditionally, the effects of increasing biodiversity on ecosystem functions are related to selection and complementarity effects (Loreau & Hector 2001; Huang at al., 2018). By quantifying how these effects shift under drought versus non-drought conditions, we uncovered the mechanisms through which diversity and spatial heterogeneity confer protection against droughts. Our results show that selection effects diminish with increasing diversity and spatial heterogeneity. In spatially heterogeneous, species-rich forests, local neighborhoods are more likely to include one or more drought-tolerant species, which are not necessarily dominant species in the absence of drought. Because drought differentially impacts tree growth (Camarero et al., 2018; Zuidema et al., 2025), these drought-tolerant species experience less growth reduction (McGregor et al., 2016; Guillemot et al., 2022), which reshapes competitive hierarchies and interaction patterns. The outcome is a more balanced performance among species, thereby weakening selection effects that are driven by dominant species.

Conversely, complementarity effects intensify with increasing diversity and spatial heterogeneity. Diverse local neighborhoods are more likely to host species with distinct ecological strategies for coping with consecutive drought events. When water becomes scarce, these strategic divergences, ranging from rooting depth to water-uptake efficiency and stomatal regulation, enable more complete partitioning of limited resources, reducing direct competition (Kröber & Bruelheide, 2014; Forrester & Bauhus, 2016; Forrester et al., 2017). Beyond resource partitioning, facilitative interactions can further ease water stress through microclimate amelioration by reducing water demand (Kunz et al., 2019; Fichtner et al., 2020). Consequently, drought-sensitive species benefit in diverse neighborhoods, where reduced competition and enhanced facilitation provide shelter from stressful environments and, in some cases, even protection from mortality. Collectively, these processes explain the amplified complementarity effects observed after consecutive droughts. Interestingly, the strengthening of complementarity effects may also arise from dynamic shifts in species interactions under stressful conditions, where competition weakens and facilitation intensifies, consistent with the stress-gradient hypothesis (Bertness & Callaway, 1994). Together, these findings demonstrate that diversity and spatial heterogeneity buffer drought effects through simultaneously equalizing species performance and enhancing complementary resource use and weaker competition or/and facilitation.

### Towards designing climate-adaptive forests

While our results provide strong evidence for positive diversity effects on drought mitigation, they reflect a limited range of functional diversity represented by the eight studied species. To reconcile the variable effects of diversity on drought responses, it is imperative to consider the broad spectrum of drought-response strategies among tree species, from strictly drought-avoiding to highly drought-tolerant types (Kramer, 1988). When interacting species share very similar functional traits, their ecological niche overlap (Rosenfeld, 2002) may cause diverse communities to respond to drought much like monocultures, thereby masking potential diversity benefits. Moreover, rising temperature under climate (IPCC AR6 (2021)) adds further complexity, as warming can promote growth under favorable conditions (Wang et al., 2023) but magnify drought stress through increased atmospheric water demand via vapor pressure deficit (Will et al., 2013; Gauthey et al., 2024). Given that species exhibit different sensitivities to these compound effects (Schillerberg & Tian 2024), rising temperatures may reshape competitive dynamics and species interactions, potentially altering the direction and magnitude of diversity effects on drought mitigation. Therefore, characterizing the functional diversity underlying drought responses is an essential next step to generalize our findings across geographic regions. Integrating multiple climate change-induced stressors into this framework will enable a more comprehensive understanding of how diversity and spatial heterogeneity can be harnessed to design climate-adaptive forests.

### Conclusions

Our results provide concrete pathways to enhance the carbon balance of human land use. Specifically, we show that both forest productivity and drought resilience are improved by tree diversity and spatially heterogeneous planting. Monoculture plantations, which currently cover roughly 140 million hectares globally (Liu et al., 2018), present a major opportunity for improvement. These forests typically sequester 2–6 t C ha^−1^ yr^−1^ in European spruce or pine plantations and 6–10 t C ha^−1^ yr^−1^ in subtropical or tropical plantations of poplar, acacia, or eucalyptus (Nabuurs et al., 2013; Poorter et al., 2016). Our data suggest that eight-species mixtures of trees can increase productivity by 11.4% under constant conditions and 41.2% under drought-disturbance regimes irrespective of spatial arrangements of the forest stand (supplement S3 and S7). Moreover, randomizing species configurations further enhances productivity, with our simulations showing 13.7% and 42.5% (supplement S3) higher biomass than block designs under constant and drought conditions in eight species mixtures, respectively. Extrapolating these results suggests that the 2–10 t C ha^−1^ yr^−1^ currently stored by monocultures could potentially be increased by 18.8%, corresponding to 2.4–12 t C ha^−1^ yr^−1^, which across 140 million hectares could sequester an additional 0.05–0.26 Gt C yr^−1^. Interestingly, the single line design is associated with relatively similar results and could constitute a good compromise between ease of management and forest sustainability. Although these are rough estimates, they illustrate that increasing forest diversity and optimizing planting design offers a strong strategy to both enhance carbon sequestration and buffer forests against drought.

## Method

We simulated tree biomass growth under drought using empirical estimates of species-specific growth rates, pairwise intra- and interspecific interaction strengths, and drought tolerance. Drought effects were imposed as species-specific growth reductions beginning in year three for three consecutive years, with recovery rates modeled using exponential functions parameterized by species-specific drought tolerance rankings. Simulations were conducted in a full-factorial design across a gradient of tree species richness (one to eight species) and four spatial planting designs: random, blocks, mini-blocks, and lines. Each drought simulation was compared to a corresponding control simulation without drought disturbance.

### Creating spatial designs along a diversity gradient

The eight species for which we estimated pairwise interactions and associated parameters served as the local species pool. From this pool, we created species mixtures at three diversity levels: 2-, 4-, and 8-species. For the 2-species mixtures, we included all possible pairwise permutations (56 unique mixtures). For the 4- and 8-species mixtures, we randomly selected 700 permutations from all possible permutations at each diversity level. We also created five replicate monocultures for each of the eight species (40 monoculture plots total) with varying initial biomass densities. This resulted in a biodiversity-ecosystem functioning (BEF) experiment comprising 1,496 plots spanning the full diversity gradient from monocultures to 8-species mixtures.

Each plot consisted of a 16 × 16 grid of trees (256 individuals total) planted at one-meter spacing, comparable to the design of the field experiment. We applied four different spatial arrangements to all mixture plots (2-, 4-, and 8-species): blocks, mini-blocks, single-line, and random designs (see Supplement S6). Monocultures had no spatial variation by definition. These four spatial designs differ in their spatial heterogeneity, which we quantified as the deviation of observed conspecific neighbors from those predicted under a null model of random spatial distribution in an 8-species mixture (Beugnon et al., 2025). Spatial heterogeneity increases progressively from blocks through mini-blocks and single-line to random designs (Beugnon et al., 2025).

### Simulating tree growth without drought events

We simulated tree growth over 10 years under varying spatial arrangements following Eq. (1) (Yu et al., 2024). This model was parameterized using Bayesian inference applied to tree growth data collected over seven years (2009–2016) in the BEF-China experiment. The fitted model integrated two components: a species-specific intrinsic growth term and a pairwise interaction term quantifying the effect of each of the eight neighboring trees j on a focal tree i. Individual tree growth was predicted as:

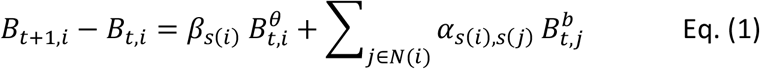

Where *B*_*t*,*i*_ is the biomass of individual tree i at time t, and *β*_*s*(*i*)_ and *α*_*s*(*i*),*s*(*j*)_ are species specific coefficients scaling the intrinsic growth and the pairwise tree interaction, respectively. The allometric scaling exponents θ and b allow for sublinear growth effects as predicted by metabolic theory.

Using this parameterized model, we simulated individual tree biomass for each forest plot arrangement (i.e., each combination of species composition and spatial design). All trees were initialized with a starting biomass of 10 grams, and growth was simulated over a 10-year period. Because all model parameters were estimated from empirical growth data spanning seven years at the BEF-China site, Eq. (1) provides a robust method for forecasting tree biomass trajectories under varying spatial arrangements at the same experimental site.

### Simulating tree growth under consecutive drought events

We simulated drought effects by reducing species’ intrinsic growth rates according to Eq. (2)

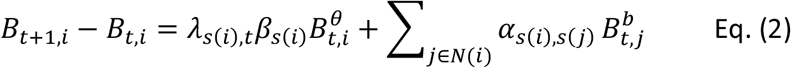

Building on Eq. (1), Eq. (2) incorporates a species-specific drought coefficient *λ*_*s*(*i*),*t*_ that quantifies the growth reduction during droughts and the years after (therefore, we have *λ*_*s*(*i*),*t*_ = 1 during pre-drought years). Its value is derived from *λ*_*s*(*i*)_ representing the growth reduction during droughts. It is based on empirical drought tolerance rankings from Fichtner et al. (2020), who measured the 50% water deficit pressure for all eight species in our study. *λ*_*s*(*i*)_ values were first sampled from a uniform distribution (0.001–0.2), and then ranked from most to least drought-tolerant using these measurements, with higher *λ*_*s*(*i*)_ values assigned to more drought-tolerant species. This approach ensures that drought-tolerant species experience smaller proportional growth reductions under water stress.

We imposed three consecutive drought events beginning in year three of the simulation. This timing reflects the heightened vulnerability of young forest stands to drought, which poses greater risks to the economic viability of restoration projects (Lalor et al., 2023).

Different species employ divergent ecological strategies and therefore exhibit varying recovery rates following drought (Hesse et al., 2023; Tao et al., 2024). We thus modeled post-drought recovery for each species using an exponential function Eq. (3):

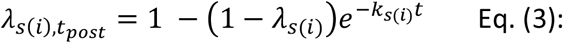

where t denotes the number of years since drought cessation. Therefore, we have 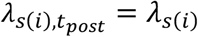 when *t = 0* (i.e., corresponding to the drought years). *k*_*s*(*i*)_ controls the recovery rate for species s(i). The parameter *k*_*s*(*i*)_ was sampled from a uniform distribution (0.01, 2.0) for each species; values approaching 2.0 allow species to recover to >90% of their pre-drought growth rate right after the drought events. Overall, this approach captures the dynamic transition from drought-reduced growth to post-drought recovery, with recovery rates varying by species according to their ecological strategies.

While species’ drought tolerance is primarily determined by hydraulic traits (Rowland et al., 2015, 2021), their recovery capacity following drought is more complex and shaped by multiple factors, including their position along the fast–slow continuum, carbon allocation strategies, and the ability to rebuild canopy and root biomass (Oram et al., 2023; Zlobin et al., 2024). Here, we assume that tree species’ recovery rate is primarily governed by their position on the fast–slow spectrum (Díaz et al., 2016). Following Fichtner et al. (2017), we used four key traits: specific leaf area, leaf nitrogen content, leaf toughness, and wood density to characterize each species’ ecological strategy. These traits are associated with productivity and shade tolerance and thus capture trade-offs between acquisitive and conservative strategies (Wright et al., 2007; Poorter et al., 2008; Lasky et al., 2014). Because conservative species invest more in durable tissues and defense at the expense of growth, we assigned the lowest recovery rate to the only conservative species. For acquisitive species, recovery rates were ranked from slow to fast according to their empirical growth rates (Fichtner et al., 2017).

Sampling of growth reduction: *λ*_*s*(*i*)_ and recovery rate parameters: *k*_*s*(*i*)_ was performed 200 times, resulting in 200 replicated BEF experiments for each spatial design. Across all diversity levels (monocultures through 8-species mixtures), spatial designs (blocks, mini-blocks, single-line, and random), and parameter samplings, we simulated biomass dynamics of a total of 1,196,800 forest plots under drought events. For each simulated plot, we recorded two response variables at the year 10: total aboveground biomass and cumulative tree mortality. Consistently with the simulations without drought events, all trees were initialized with a starting biomass of 10 grams, and growth was simulated over a 10-year period.

### Calculating selection and complementarity effects

In order to gain a mechanistic understanding of how diversity and spatial heterogeneity buffer drought effects, we applied an additive partitioning approach (Eq. (4); Loreau & Hector, 2001) quantifying complementarity effects (CE) and selection effects (SE) under drought versus non-drought. This analysis required recording the total biomass of each species per plot at year 10 under both drought and non-drought conditions.

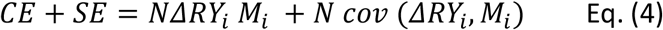

with N being the diversity of the mixture, *NΔRY*_*i*_ captures the deviation of the observed from the expected relative productivity of species i. *M*_*i*_ denotes monoculture productivity of species i. Complementarity effects quantify the average change in productivity of species in mixture compared to their monoculture performance, reflecting niche partitioning and facilitation. Selection effects quantify the covariance between species’ monoculture productivity and their success in mixture, capturing whether mixtures are dominated by species that perform better or worse than the average monoculture. Because the additive partitioning approach requires monoculture biomass data for all component species in a mixture, we excluded simulation replicates in which drought events caused complete mortality in one or more monocultures. All simulations, calculations, and data visualizations were performed using R software version 4.3.3 (R Core Team, 2024).

## Acknowledgements

We acknowledge the support of the BEF-China platform. WY was supported by the International Research Training Group TreeDì funded by the German Research Foundation (DFG; GRK 2324, 319936945). GA and BR gratefully acknowledge the support of iDiv funded by the German Research Foundation (DFG–FZT 118, 202548816).

